# Data-driven modeling of leading-following behavior in Bechstein’s bats

**DOI:** 10.1101/843938

**Authors:** Pavlin Mavrodiev, Daniela Fleischmann, Gerald Kerth, Frank Schweitzer

## Abstract

Leading-following behaviour in Bechstein’s bats transfers information about suitable roost sites from experienced to inexperienced individuals, and thus ensures communal roosting. We analyze 9 empirical data sets about individualized leading-following (L/F) events, to infer rules that likely determine the formation of L/F pairs. To test these rules, we propose five models that differ regarding the empirical information taken into account to form L/F pairs: activity of a bat in exploring possible roosts, tendency to lead and to follow. The comparison with empirical data was done by constructing social networks from the observed L/F events, on which centralities were calculated to quantify the importance of individuals in these L/F networks. The centralities from the empirical network are then compared for statistical differences with the model-generated centralities obtained from 10^5^ model realizations. We find that two models perform well in comparison with the empirical data: One model assumes an individual tendency to lead, but chooses followers at random. The other model assumes an individual tendency to follow and chooses leaders according to their overall activity. We note that neither individual preferences for specific individuals, nor other influences such as kinship or reciprocity, are taken into account to reproduce the empirical findings.

## 1 Introduction

In most social species, individuals need to coordinate their actions in order to maintain group cohesion and to obtain grouping benefits such as improved foraging, energetic savings from huddling, or a reduced predation risk (Krause and Ruxton, 2002; Conradt and Roper, 2005; Sumpter, 2005; Kerth, 2010). Typically, such coordination is achieved via information transfer from informed to uninformed (naive) individuals, as in the case of collective motion in fish swarms and social insects (Franks *et al*., 2002; Seeley *et al*., 2006; Ward *et al*., 2011).

In this paper, we focus on information transfer in maternity colonies of Bechstein’s bats (*Myotis bechsteinii*). This bat has several key traits that make it particularly relevant for studying coordination problems in animal systems (Kerth and Reckardt, 2003; Kerth *et al*., 2006). First, the frequent switching of communal day roosts implies that group coordination and collective decision-making regarding communal roosting is vital for individuals getting grouping benefits. Second, coordination must be achieved in the presence of different individual preferences and limited information about the suitability of roosts (Fleischmann *et al*., 2013), which has been recognized as particularly challenging for collective decision-making (Conradt, 2011)

As Bechstein’s bats forage solitarily (Melber *et al*., 2013), they have only limited information of the nightly exploration behaviour of others. Moreover, some individuals are more active in exploring their habitat and thus become better informed about the location of suitable roosts than their less active colony mates (Kerth and Reckardt, 2003; Fleischmann *et al*., 2013). Information asymmetry and heterogeneous roosting preferences in the colony are, thus, invariable challenges in coordinating roosting behaviour and ensuring grouping benefits.

All these challenges not only make it difficult for the colony members to achieve group coordination, they also make it hard for us to infer which influence, on the individual level, is relevant for the outcome. The information transfer from informed to naive individuals does not explain how such individuals find each other. Recruitment plays an important role (Richner and Heeb, 1996; Kerth and Reckardt, 2003), but the rules by which individuals *form leading-following pairs* in which an informed leader guides an uninformed follower to a new roost, still remain unknown. Hence, questions whether some individuals exert disproportionate influence by assuming more or less fixed leadership roles or whether followers randomly select leaders cannot be answered.

Some animal studies have suggested that leadership is a personality trait independent of differences in information or knowledge of the environment (see Johnstone and Manica (2011) and references therein). However, detecting this personality trait becomes complicated when observations do not continuously track individuals, but rather contain isolated measurements, e.g. discrete records of animal occurrences at measurement sites, as it is the case of roost monitoring in the studied Bechstein’s bat colonies. As a consequence identifying influential individuals or the emergence of distinct roles is contingent on our ability to reconstruct reliably the missing information on inter-individual interactions. This problem is addressed in a forthcoming paper (Mavrodiev *et al*., 2019) which proposes a methodology to infer leading-following (L/F) events in Bechstein’s bats from such discrete records of occurrences.

In this paper, we build on data sets about these L/F events from two different colonies and up to five years (see Section 2.1 for details). We demonstrate how we can infer the rules underlying the formation of L/F pairs from this data, by introducing and testing five null-models that vary in complexity regarding the information involved.

Social network theory plays an important role in our approach. First of all, we represent the interaction between individuals as a social network that contains all L/F events. The social networks constructed from the L/F events (Mavrodiev *et al*., 2019) occurs on a time scale much shorter than the long-term social relationships detected in Bechstein’s bats (Kerth *et al*., 2011).

Secondly, we use the L/F networks to quantify the importance of individuals in leading and following by means of a centrality measure. Our models contain different hypotheses about the formation of such L/F networks. Specifically, we consider random influences, observed activities, and individual tendencies to either lead or follow in the formation of L/F pairs. We test the performance of our models by comparing the predicted individual centralities with the ones obtained from the empirical network. The results allow us to infer possible sets of rules that may lead to the observed recruitment events, and to deduce from these rules the impact of different information in forming L/F pairs.

## 2 Materials and methods

### 2.1 Study animals and data

In this paper we analyze data about leading-following events in Bechstein’s bats, a behavior that facilitates communal roosting. This bat species forms colonies of about 10-50 individuals where females communally give birth and nurse and wean their offspring. For their daily roosting, e.g. in tree cavities and bat boxes (Kerth and König, 1999; Kerth *et al*., 2011), they form one to several roosting groups that occupy distinct day roosts to benefit from grouping, e.g. via social thermoregulation (Pretzlaff *et al*., 2010). The formation of roosting groups requires the coordination of decisions about where to roost.

Field experiments have shown that, to achieve this critical coordination, Bechstein’s bats engage in information transfer via *leading-following behavior* (Kerth and Reckardt, 2003). During their nightly habitat exploration they accumulate private information about the location of potential novel roosts. Experienced individuals then transfer this private knowledge to naive conspecifics by leading them to these locations. This defines *leading-following* (L/F) events characterized by a leading individual who *recruits* a follower and leads it to a particular roost. However, the inter-individual rules governing recruitment are largely unknown. Hence, this paper aims at revealing such rules from the observed L/F events.

We build on extensive longitudinal data sets from two Bechstein’s bat colonies, GB2 and BS, for which bat movements in and out of experimental roosts have been recorded. For these recordings bats have been marked with individual RFID-tags. This assigns a unique 10-digit ID to each bat that allows to identify its entry in specially prepared bat boxes, using automatic reading devices. `#readings` in Table 1 gives the total number of such recordings for each colony and year. These recordings were pre-processed by specific algorithms that are able to detect L/F events and to distinguish them from swarming behaviour (Mavrodiev *et al*., 2019). As the result, we are provided with tables of consecutive L/F events (for an example see Table S1 in the Supplementary Material), which are used as an input for our study. Table 1 gives an overview of the data sets used in this paper.

**Table 1:** Basic structural properties of the leading-following networks from the GB2 and BS colonies. Shown are number of nodes (bats), number of identified L/F events (links) for different years. The data was inferred from collected raw data as described in (Mavrodiev *et al*., 2019).

| Colony | Year | #bats | #readings | #L/F events |
| --- | --- | --- | --- | --- |
| GB2 | 2008 | 34 | 4243 | 262 |
|  | 2011 | 16 | 1929 | 86 |
| BS | 2007 | 16 | 5600 | 169 |
|  | 2009 | 17 | 9102 | 201 |
|  | 2010 | 19 | 2169 | 148 |
|  | 2011 | 7 | 2016 | 26 |

### 2.2 Individual activities

A second look at the data set about L/F events already tells us that individuals are not equally participating in discovering roost boxes and in forming leading-following pairs. To capture this heterogeneity, we distinguish between three different empirical measures.

Activity *a*^*i*^ of an individual *i* is defined as the total number of readings that involved this particular bat *i* during the study period. This captures also visits of boxes during the individual exploration, i.e. it includes discovery and revisits that are unrelated to leading-following behaviour. In fact, as the comparison of `#readings` and `#L/F events` in Table 1 shows, most readings came from discovery, exploration, and revisits. In Figure 1 we plot the activity *a*^*i*^ of all bats of the colony GB2 in the year 2008. To make differences more visible, the bats are ranked according to their activity *a*^*i*^. To be comparable to other colonies and other years, we always normalize *a*^*i*^ to the total number of readings given in Table 1.

**Figure 1:**
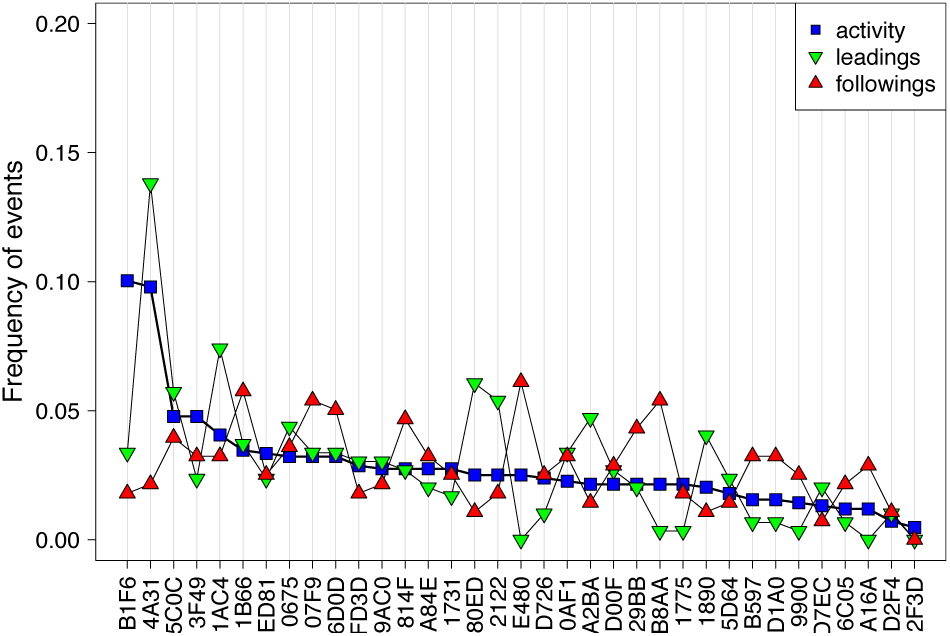
Normalized numbers of readings (activity), leading and following events for each individual of the colony GB2 in 2008. Ranking according to nurmalized activity.

The other two empirical measures capture the involvements of bats in leading-following events. *l*^*i*^ (*f*^*i*^) gives the number of L/F events a given bat *i* was involved as a *leader (follower*), normalized to the total number of L/F events given in Table 1. Different from the general activity in exploring boxes, these two values describe the “activity” in the information transfer as a result of the recruitment process. We note that these measures do not tell us, which individual initiated the formation of a *leading-following* (L/F) *pair*. It could be the follower, actively seeking for a leader or the other way round.

In Figure 1 we also plot the normalized values *l*^*i*^ and *f*^*i*^ for each individual. We argue that these values represent an *individual tendency* to either lead or follow, which may have its reasons in other individual characteristics, such as age, weight, reproductive status, etc. Here, we take them into account as empirical facts. As the plots in Figure 1 show, only a small number of individuals stands out regarding their activity. Also, we already note that leading and following events are not trivially related to activity. In fact, to explore this relationship is the main goal of our null models, which will be further discussed below.

In a next step, we construct from the L/F events a social network in which nodes represent bats and directed links represent individual L/F events. A link *A → B* means that the naive individual *A* follows the experienced individual *B*, hence information is transferred from *B* to *A*. We aggregate over the time interval of one season (i.e. one year), to obtain *weighted and directed L/F* networks, in which the weights of the links are defined by the frequency of the respective events.

Two examples of the constructed L/F networks are shown in Figure 2. They refer to the same colony, BS, but to different years, 2007 and 2011. We observe that the size and the composition of the colony has changed drastically, due to external influences and birth/death processes (Baigger *et al*., 2013; Fleischer *et al*., 2017). Table 1 gives an overview of the size of the L/F networks for the different years.

**Figure 2:**
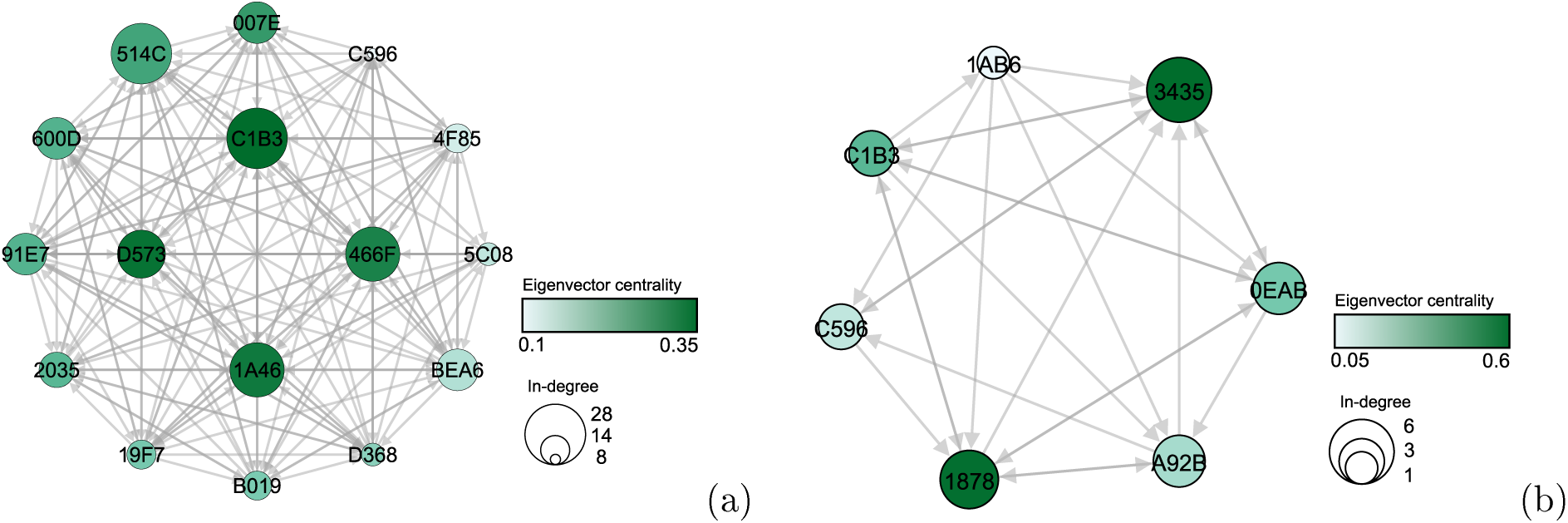
Aggregated leading-following network for the BS colony in (a) 2007 and (b) 2011. Individual bats are represented as nodes and identified by the last 4 digits of their RFID tag. Directed edges represent following behaviour. Node colors indicate eigenvector centrality (see main text), whereas node sizes indicate in-degree centrality. The total number of L/F events, including multiple leading-following between the same individuals, is (a) 290 and (b) 52. The total number of unique L/F events is (a) 169 and (b) 26 (Table 1). Note that for the sake of illustration, edges show only *unique* L/F events, i.e. leading-following between the same leader and follower, but to different roosts, are omitted.

We can now use the aggregated L/F network to quantify the importance of an individual, i.e. of node *i*, in the transfer of information. For this, we build on established measures, in-degree centrality, 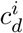, and eigenvector centrality, 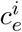 (see Mavrodiev *et al*. (2019) for details). In-degree centrality only considers the number of unique followers, i.e. the number of incoming links (denoted as in-degree). In the given weighted network, 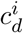 also considers the weights, i.e. the frequency of following events. It is a measure of *direct influence* between individuals.

However, as the transfer of information can also happen through intermediate individuals we need to consider *indirect influence* in addition to the direct one. Indirect influence is captured by the eigenvector centrality that considers the centrality of the followers. An experienced bat that leads a few bats that themselves lead other bats has a higher eigenvector centrality, i.e. it is more influential in the transfer of information than a bat that leads many other bats that never lead.

Eigenvector centrality of an individual grows with the length of the chain of direct and indirect followers. Hence, individuals that are part of a longer chain are automatically considered more influential than individuals leading many followers. This is considered a drawback for the given application. Therefore, we have corrected for this by introducing a new metric, *second-degree centrality*, 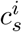, that combines the above mentioned centralities using a weight *β*:

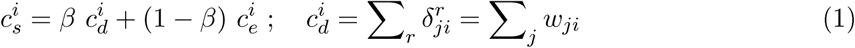

Here *δ*_*ji*_ denotes an L/F event, i.e. *δ*_*ji*_ = 1 every time *j* followed *i*, and zero otherwise. The summation *r* goes over all detected L/F events. The sum over such L/F events, with respect to a given bat *j*, equals the weight, or the frequency, *w*_*ji*_, and the sum over all bats gives the total weighed in-degree. For the eigenvector centrality 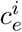 we cannot give a closed form expression because it is obtained from solving the eigenvector problem *λ* ⋅ **c**_*e*_ = *A* ⋅ **c**_*e*_, where **c**_*e*_ is the eigenvector, *A* is the adjacency matrix that contains information about the directed and weighed links, and *λ* is a scaling factor (for an example of how to calculate the eigenvector, see e.g. (Mavrodiev *et al*., 2019)).

Figure 2 illustrates how the different centralities vary across individuals. In the example of the (rather small) colony BS, we observe that every individual was involved in L/F events, hence the network consists of only *one* strongly connected component (SCC), i.e. there is a path from any node to any other node via the directed network. Because the network is rather dense, the individual in-degree centralities are not very different, however the eigenvector centralities are, as indicated by the color scheme. Hence, it makes sense to combine this information in the *second-degree centrality*, 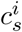, as explained above. It computes importance from the in-degree of the focal individual and the sum of the in-degrees of its followers (i.e. the followers of the followers, or second-nearest neighbors), weighted by the factor *β*. In the following, we choose *β* = 0.5 and use second-degree centrality to quantify the influence of individuals in information transfer.

### 2.3 Model testing

The methodology outlined above allows us to assign an importance value 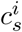 to every individual based on the L/F network that was reconstructed from the observational data. In the following, we consider these centralities 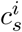 as *ground truth*, because they have been directly calculated from the empirical data.

At this point we face the problem that the L/F network contains important information about the interaction between bats. However, it does not tell us anything about how this interaction came into place. I.e., we miss the rules by which individuals form leading-following pairs that are later discovered in the data. To find out about possible rules that are compatible with the ground truth, we will test different sets of rules, called models, that may govern this process. For each of these models, indicated by the index *m*, we calculate the resulting importance values 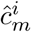 and compare them with the respective values 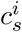 from the ground truth. We deem a model successful if it is able to reproduce the ground truth statistically.

Specifically, we run each model 10^5^ times and compute from this the distribution 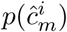 of each individual importance 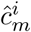. From this distribution, we estimate its *Gaussian kernel density*, which is a function 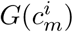. The Gaussian kernel density evaluated at the empirical centrality, 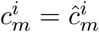, defines an *individual density score*, 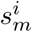. This score indicates to which extent the observed influence, 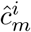, of an individual is reproduced by the given model *m*. Choosing this method, we account not only for the presence of the 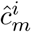 in the *95% confidence range*, but also for their *likelihood* within the individual distributions 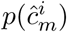.

To evaluate the overall performance of a given model *m*, we sum up the individual density scores 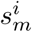 to produce the *total density score*, 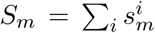. The larger the *S*_*m*_, the better the given model reproduces our empirical finding of the individual centralities 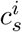.

### 2.4 Model generation

It remains to specify how the different models should be generated. Our models reflect rules for the interactions between two individuals. Specifically, they tell how, i.e. based on which information, a *leader* and a *follower* form a leading-following pair. Since such rules are not known a-priori for an animal system, we present a method of *incremental* null-model building, where each null model is based on a hypothesis about these rules. That means, we start our model-building from null models with very simple hypotheses and incrementally add rules of increasing complexity. The calculated total density score, *S*_*m*_, tells us whether the addition of new rules has improved the quality of the model in comparison with models already tested.

Null models are recognized as useful in the presence of inherently non-independent behavioural data (Farine, 2017; Croft *et al*., 2011). They serves as a sieve that can help us reveal the minimum complexity required to reproduce the observed individual influence. Importantly, in addition to suggesting the most parsimonious recruitment mechanisms, null models serve to focus research attention by identifying those cases, in which a more complex mechanism may be at play. This is particularly useful in biological systems, in which high individual diversity inherent to the system makes it harder to identify common determinants of observed behaviour.

For models with comparable score, *S*_*m*_, we follow Occam’s razor and deem the model with the least assumptions as the better one. Complexity in our case is expressed by the information that is needed to apply certain rules. In leading-following behaviour of Bechstein’s bats this information can include past experience, kinship, reproductive status, age, etc. However, as our models start with minimal complexity, we consider only information that is already available from the data. We remind that empirical evidence is provided, at the aggregated level of the individual, for the activities, *a*^*i*^, and for the frequencies of being a leader, *l*^*i*^, or a follower, *f*^*i*^. Additionally, we also have, from the L/F events themselves, detailed information which individuals *i* and *j* are involved in forming a leading-following pair.

This empirical information also sets *boundary conditions* for possible models. Specifically, if we want to compare the model outcome, expressed by the centrality values 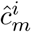, with the empirical values 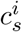, we have to use the same *aggregated* network properties. That means, we have to take the empirical numbers of individuals and of L/F events as constraints for our models. These values are given in Table 1. We emphasize that these are *aggregated* values, i.e. by fixing the numbers of individuals and of L/F events we basically keep the *density* of the resulting L/F network constant. But we do *not* fix the topology, i.e. the detailed structure of the links between nodes, by respecting these boundary conditions.

With a fixed number of nodes, i.e. bats, and a fixed number of directed links, i.e. L/F events, our models can be seen as rules to *rewire* a given L/F network. Rewiring means that we create a leading-following pair based on certain *rewiring rules* that take different information into account. In the simplest possible manner, we can, for example, create pairs between randomly chosen leaders and randomly chosen followers such that the given number of L/F events is respected. This would be one example for a rewiring rule that *does not* take any information into account.

More complex rewiring rules, however, should consider the aggregated information given by the available empirical data, specifically *the three information* about *activity*, *a*^*i*^, *tendency to lead*, *l*^*i*^, and *tendency to follow*, *f*^*i*^. Hence, in the following, we will discuss models in which the rewiring rules always specify to what extent this information is considered. That means, each model consists of a *hypothesis* about the role of the three information in forming leading-following pairs. If such a model is able to describe the observed centralities better than another model, we argue that such information play an important role to form the respecting leading-following pairs. In the spirit of parsimony, we contend that the hypothesis underlying this model is a tenable candidate for the *real mechanisms* underlying leading-following behaviour. But, of course, we cannot exclude that other possible hypotheses, based on the use of other information, can describe the empirical findings as well.

## 3 Results

### 3.1 Evaluating models with different complexity

Below we present five null models, together with their underlying hypotheses about rules to form leading-following pairs. We start with the simplest model, already mentioned above, which shall serve as a reference to judge the impact of more complex rules, afterwards.

#### [M1] Random behaviour

This model assumes that *none* of the above mentioned information about individual activity, tendencies to lead or to follow is taken into account. Consequently, for every leading-following pair, we choose *both* the leader and the follower *randomly*. For a network consisting of *n* nodes, this means that every node has a equal probability of 1*/n* to be chosen as a leader or a follower. We note again that this way only the total number of L/F events is considered, but no information about the composition of L/F pairs.

The formation of L/F pairs according to these rules occurs until the total number of L/F events known from empirics is reached. We then take the L/F network formed this way, to calculate the corresponding centrality values 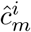 of each node. In order to obtain the distribution 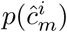, we repeat this procedure 10^5^ times. Afterwards we calculate the individual density scores 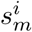 as explained above.

The results for 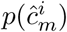 are shown in Figure 3 for our running example, the colony GB2 in year 2008, for all other data sets we present the corresponding plots in the Supplementary Material. The *x−*axis lists all individuals of the colony, identified by their abbreviated RFID tag ID. The *y−*axis displays the second-degree centrality of each individual as obtained from the empirical L/F network. The centrality scores are always normalized with respect to the largest centrality score (empirical or model-generated) in the corresponding model. Precisely, if the empirical score is higher, the centrality values are normalized such that the largest empirical value equals one. However, if the model-generated score is higher then, because of the normalization of the centrality values, the largest *empirical* value is less than one.

**Figure 3:**
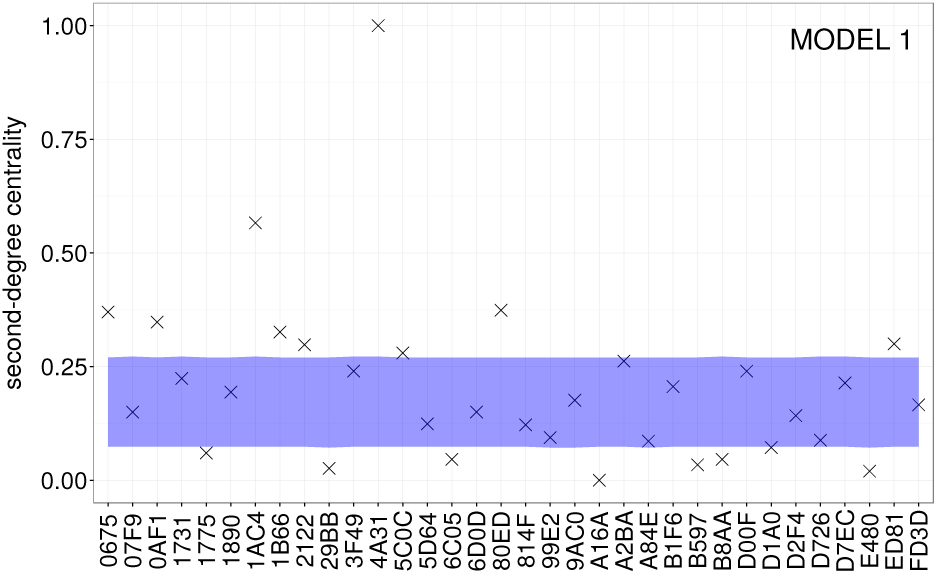
Model-generated vs. empirical scores for colony GB2 in 2008, using [M1].

As explained before, the values indicated by the (x) symbols serve as the ground truth for evaluating the model. We observe that the centralities are quite heterogeneously distributed. For a majority of individuals, we find low to mid values, only a minority has high importance values.

The blue band in Figure 3 denotes the 95% inner-most range of the model-generated centrality distributions, 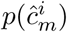. This range represents our model-specific expectation for the centralities of each individual, given the information about the respective colony members used to form the L/F pairs. The mismatch with the empirical data is obvious, but not surprising. If there was a match, this would imply that bats form L/F pairs entirely at random, not taking any information into account. We can now observe how the model prediction improves if we test gradually more complex rules for forming L/F pairs.

#### [M2] Random behavior and activity

This model assumes that active bats are more likely to be selected as leaders than less active individuals. The rationale for this assumption is that more active bats have a higher likelihood to be followed, as their frequent flights render them more “detectable” to potential followers. The normalized activity, *a*^*i*^, of a bat was defined in Section 2.2 and now determines the probability of being chosen as a *leader*. Followers are still chosen equally at random, as before, i.e. *no information* about the tendency to lead or to follow is taken into account to form L/F pairs.

The results are shown in Figure 4, which can be compared to Figure 3. It becomes obvious that rules which take information about individual *activities a*^*i*^ into account, perform better compared to [M1]. But still, there is a considerable mismatch between the model and the empirical data.

**Figure 4:**
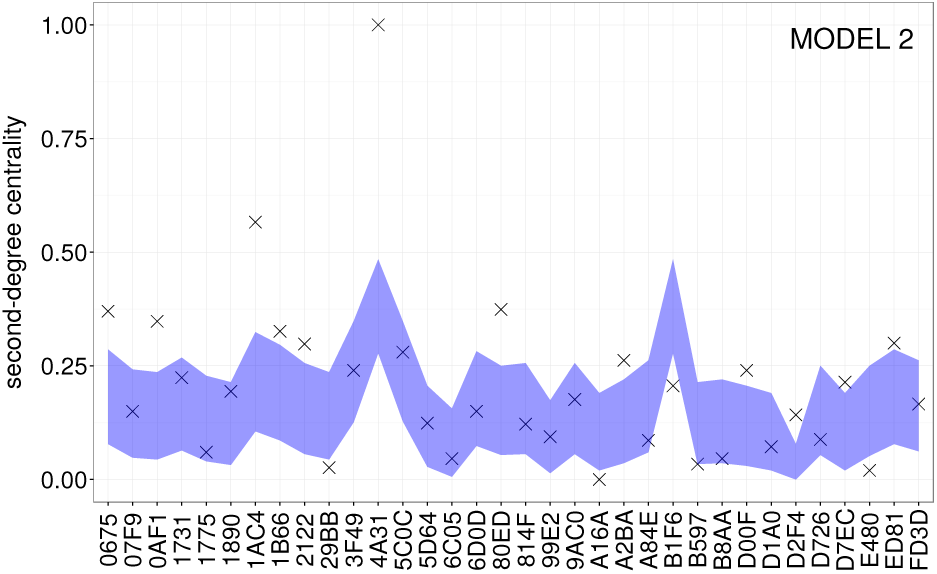
Model-generated vs. empirical scores for colony GB2 in 2008, using [M2].

#### [M3] Individual tendency to follow

This model assumes that the individual tendency to follow plays the major role in forming L/F pairs. The normalized value, *f*^*i*^, of a bat was defined in Section 2.2 and now determines the probability of being chosen as a *follower*. The probability of being chosen as a leader is still equal at random, i.e. *no information* about the tendency to lead and no information about activity is taken into account. In the implementation of the model, for each link in the L/F network we keep the follower as observed in the data and only rewire the link to a randomly chosen individual as leader.

If this model performs well, it implies that the formation of L/F pairs is driven by the followers, which would potentially follow any randomly chosen other bat. But, as the results in Figure 5 show, taking the individual tendencies to follow, *f*^*i*^, into account, gives results better than [M1], but not better than [M2].

**Figure 5:**
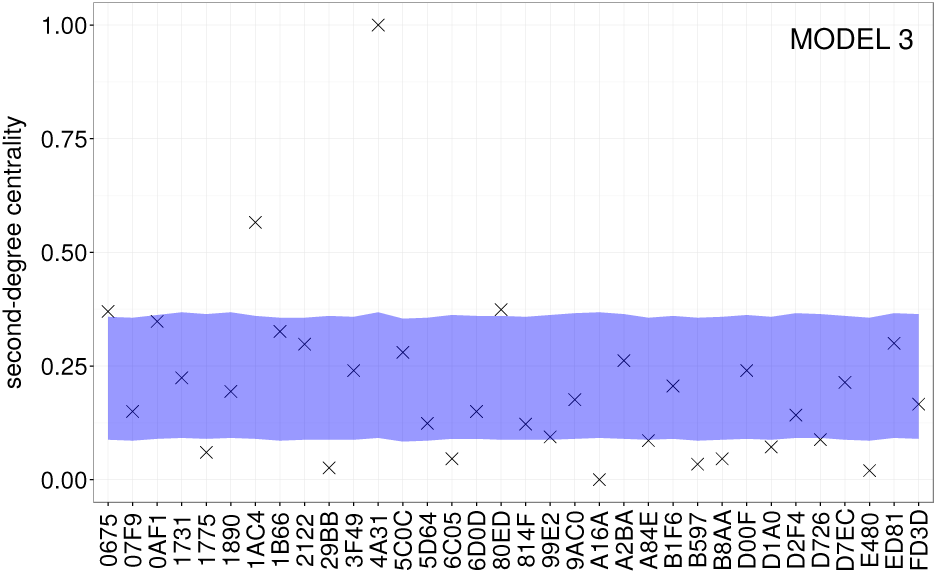
Model-generated vs. empirical scores for colony GB2 in 2008, using [M3].

### [M4] Individual tendency to lead

To be contrasted with [M3], this model assumes that the individual tendency to *lead* plays the major role in forming L/F pairs. The normalized value, *l*^*i*^, of a bat was defined in Section 2.2 and now determines the probability of being chosen as a *leader*. The probability of being chosen as a *follower* is still equal at random, i.e. *no information* about the tendency to follow and no information about activity is taken into account. In the implementation of the model, for each link in the L/F network we keep the leader as observed in the data and only rewire the link to a randomly chosen individual as follower.

If this model performs well, it implies that the formation of L/F pairs is driven by the leaders, whereas followers can be any randomly chosen other bat. The results shown in Figure 6 indeed demonstrate that taking such information into account remarkably improves the performance of the model. Not only that the 95% confidence range includes (almost) all empirical values, the range has also become very narrow, which indicates a quite precise expectation from the model. Comparing this model with [M2], we also see that information about the (quite specific) tendency to lead is more important in forming L/F pairs than information about the (quite general) activity of bats.

**Figure 6:**
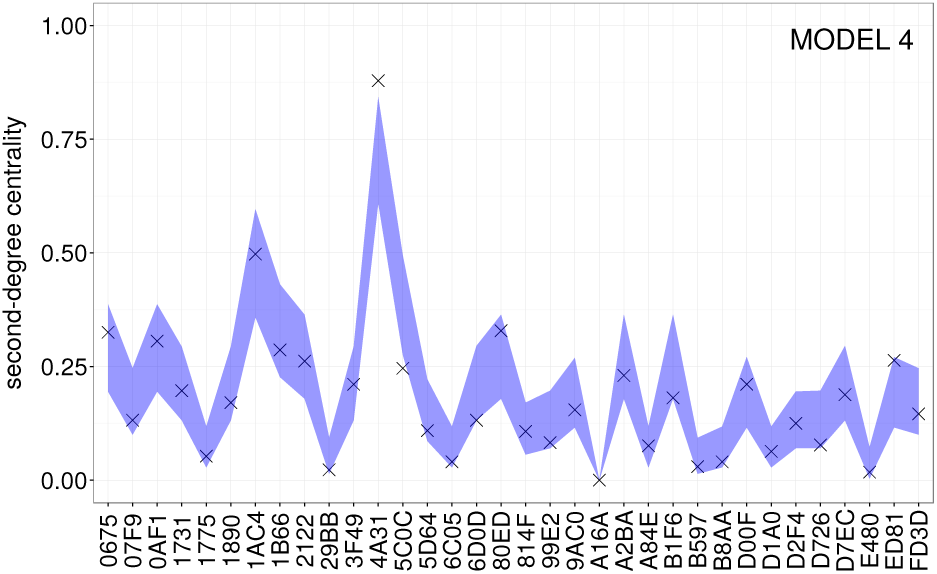
Model-generated vs. empirical scores for colony GB2 in 2008, using [M4].

#### [M5] Individual tendency to follow and activity

This model combines [M3] and [M2], i.e. it assumes that the tendency to follow, *f*^*i*^, plays a major role in forming L/F pairs. However, the leaders should not be chosen equally at random, but by taking information about their activity, *a*^*i*^, into account. Again, we argue that more active bats are more likely to be chosen as leaders. In the implementation of the model, for each link in the L/F network we keep the follower as observed in the data and only rewire the link to a leader chosen with a probability equal to the relative activity *a*^*i*^. Importantly, the association between a leader and a follower is still random, i.e. bats do not have special preference to lead or follow a specific individual.

The results are shown in Figure 7 and indicate also a good performance of the model in reproducing the empirical centrality values. Thus, we need to quantify differences in the performance of the models, below.

**Figure 7:**
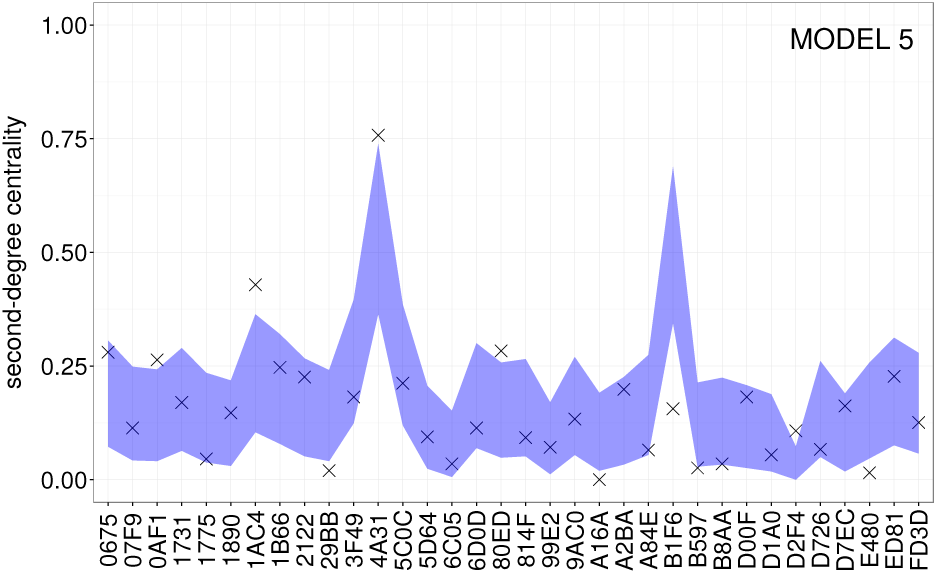
Model-generated vs. empirical scores for colony GB2 in 2008, using [M5].

In conclusion, we notice the considerable mismatch between empirics and model results for the models [M1], [M2] and [M3]. Of course, quite a large number of measured centrality values fall into the 95% confidence range, but only because this band is very broad for the models [M1], [M2] and [M3]. The model results do not allow us to deduce that the formation of L/F pairs, which is the basis of the evaluation, can be reasonably described by these models, i.e. by only taking the respective information into account. This is different for the models [M4] and [M5] that offer a significant improvement, also because their confidence bands are quite narrow, i.e. give a precise prediction. We note that [M4] takes one information into account, namely the tendency to lead, *l^i^*, whereas [M5] takes two information into account, namely the tendency to follow, *f*^*i*^, and the activity of the leader, *a*^*i*^.

### 3.2 Comparing model performance for all data sets

To further distinguish the performance of the five different models, we have to extent the above analysis to *all* data sets from all years. To quantify the performance, we will use the total density score *S*_*m*_ that was already introduced as a goodness measure in Section 2.3. Table 2 shows this comparison between the five models in terms of their total density scores.

**Table 2:**
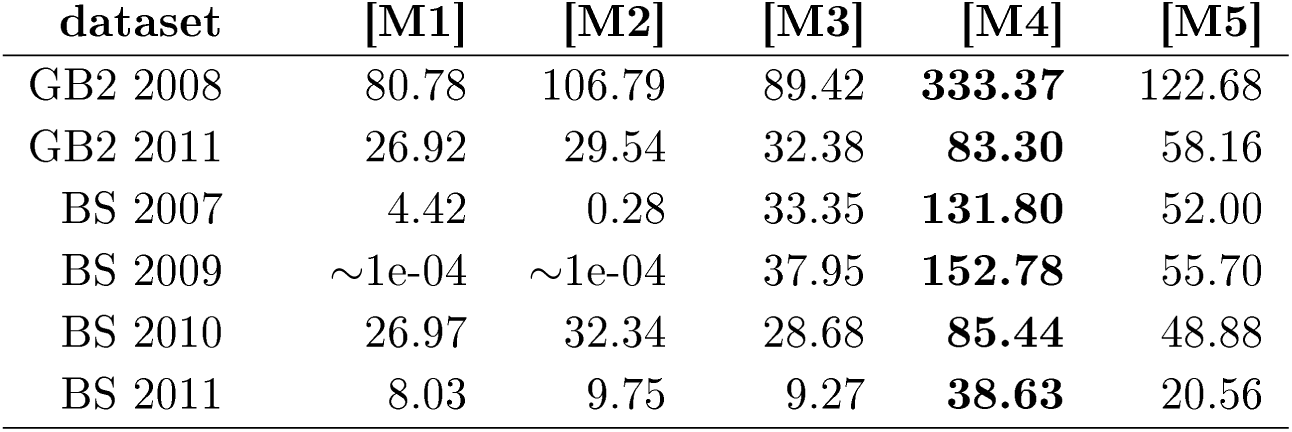
Comparison of the models [M1]-[M5] by means of their total density score *S*_*m*_ (see Section 2.3). The higher the values of *S*_*m*_, the better the model reproduces the empirical findings.

We find that model [M1] has the lowest total density score in all data sets (except for BS 2009, where it has the second-lowest value). The poor performance of [M1] is a clear indication that L/F events are not formed entirely by chance and individuals are not equally likely to be leaders or followers. Compared to [M1], models [M2] and [M3] offer a slight improvement. However, for all data sets [M2] and [M3] are outperformed by model [M5]. This shows that information about activities of leaders in [M2] or the tendency to follow in [M3], *taken separately*, cannot explain the formation of L/F pairs better than the combination of both information, as considered in [M5].

However, we find that [M4] is the model with the highest total density score in Table 2. This was already indicated by Figure 6, where the distributions of centralities generated by [M4] closely match those extracted from the field data (See also Section S.2 in the Supplementary Material). Here, we confirm that [M4] performs best among the five proposed models for all data sets from the two colonies for all years. Model [M5] consistently performs second-best among the five proposed models.

## 4 Discussion

As outlined in Farine (2017), null models are important and even indispensable tools in social network studies. When data come from non-independent observations of multiple individuals the corresponding network representation can easily generate patterns that look like social structures. Null models allow us to test informed hypotheses by restricting alleged social relationships and thereby accounting for non-social factors that may have produced the given network pattern. In the process, we are able to propose viable individual-level interaction mechanisms that then result in justified social structures on the network level. In this paper we have applied an incremental null-model building process in the context of leading-following behaviour.

### Methodological approach

Leading-following (L/F) behaviour is used to transfer information about suitable roosts in Bechstein’s bats, to ensure communal roosting (Kerth and Reckardt, 2003). However, the behavioural rules by which bats form such L/F pairs are largely unknown. Therefore, the main goal of this paper is to infer (sets of) minimal rules that comply with the observed L/F events detected in empirical recordings of box visits. This implies solving the following methodological problems.

First, we need to define how we want to compare the model outcome with a ground truth obtained from empirics. Here, we propose the comparison on the level of aggregated individual measures. Specifically, we construct a (directed and weighted) social network that contains all L/F events and calculate, based on this network, individual centralities as a measure of importance in transferring information. Hence, we build on *social network theory*, (i) to construct the L/F networks, and (ii) to quantify the positions of individuals in this network, using established centrality measures.

On the modeling side, we propose five null-models (sets of rules) for forming L/F pairs. Each model can be seen as a randomization process of the given L/F network, taking additional information as constraints into account. Running these models 10^5^ times each, we obtain an ensemble of model-generated L/F networks, from which we calculate model-generated individual centralities. These are then compared to the empirical centralities using a statistical procedure that assigns a individual density score 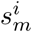 to each predicted centrality. The sum over these scores, *S*_*m*_, defines the performance of the model *m* in relation to other models tested. The higher *S*_*m*_, the better the prediction. Hence, we cannot obtain an absolute best model because we cannot test all possible models. However we can deduce which sets of rules perform comparably better in capturing the ground truth.

#### Inferring rules for forming leading-following pairs

The second methodological problem regards the generation of the null-models. Here, we follow an approach to incrementally build null-models of increasing complexity. All null-models can take the following empirical information into account: (i) individual activity, *a*^*i*^, (ii) individual tendency to lead, *l*^*i*^ and (iii) individual tendency to follow, *f*^*i*^. The sets of rules, i.e. the complexity of the models, differ in which of these information are considered for forming L/F pairs. In total we have tested five different models. If a model *m* performs better, i.e. has a higher total density score *S*_*m*_ in comparison with other models, we argue that the respective information taken into account plays an important role in the rules to form L/F pairs.

This procedure allows us to test hypotheses about information involved in recruitment processes for L/F events. By incrementally increasing the complexity of the hypotheses, we can understand behavioral patterns as the emerging result of individual interactions that, in the ideal case, make biological sense.

According to our investigations, what information do bats take into account when forming leading-following pairs? From our model testing, we can exclude that bats form L/F pairs at random. This sounds trivial, however, it needs to be tested - and the random formation serves as a reference case to understand the impact of more complex rules. We can further exclude that information about the *activity* of leaders *or* information about the tendency to *follow alone* are sufficient to explain the formation of L/F pairs. Instead, we found that a *combination* of information about activity of leaders and the tendency to follow is necessary to sufficiently explain the observed formation of L/F pairs. The respective model [M5] performs second best in capturing the empirical centralities.

The best performance, however, was obtained in model [M4] where information about the tendency to lead, *l*^*i*^, determines the formation of L/F pairs. This implies that leaders play the most important role in the recruitment process, whereas for followers no additional information is needed, i.e. they randomly select leaders. We note that such random association supports existing findings (Kerth and Reckardt, 2003) on the lack of kinship and reciprocity in the recruitment process, and model [M4] serves as additional evidence for this.

### Role of indirect influence

Because model [M4] shows the best performance, in the following, we now critically examine the so-called “threats to internal validity” of this result. The basic hypothesis underlying [M4] is that information about the tendency to lead determines the formation of L/F pairs. Hence, in our implementation we have fixed the leaders according to their empirical occurrence, i.e. we have set the tendency to lead to the empirical value, while followers are chosen at random. This is equivalent to a *fixed in-degree* of the leaders. Our comparison measure is the *second-degree centrality*, 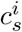, of each individual. This measure is, of course, not completely independent of the in-degree, but has to reflect its influence simply because it is calculated on the aggregated network of L/F events. But, as a centrality measure, it contains much more information, in particular about indirect followers and their importance.

An indirect follower *C* of individual *A* receives information about a possible roost not directly from *A*, who discovered the roost, but from an intermediate individual *B* who was the *follower* of *A*, and the *leader* of *C*. Such chains of information transfer *are* captured in the *second-degree centrality*, 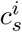. If these chains do not play a role, this could also result from the fact that they do not exist. Mavrodiev *et al*. (2019) have provided an analysis of the length of chains in L/F events. They found that even long chains can be detected, but their probability of occurrence decays exponentially. Chains of length 1, i.e. single L/F events, make 65%, while chains of length 2 make about 18% and chains of length 3 still make about 8% of the findings.

Hence, the existence of chains of information transfer along two or more individuals *cannot* be ignored. That is why it makes sense to consider second-degree centrality instead of simple indegree centrality. But the influence of chains of information transfer, due to their relatively low occurrence, is not large enough to produce relevant deviations between modelled and empirical centrality values. I.e., we can conclude that this additional information does not play an important role in forming L/F pairs.

#### Role of flight activity

The second promising model to explain the formation of L/F events is [M5]. It performs second-best in comparison to [M4], but, as we have explained, this is due to the fact that the information used in [M4] partly correlates with the measures used for comparison. Given this, we can deem [M5] a valid candidate to explain the rules to form L/F pairs.

[M5] is different from [M4] in that the rules use two different information about the leader *and* the follower, namely the activity of the leader and the tendency to follow. In this respect, the complexity of [M5] is larger. However, there is no preference of a follower to choose a *specific* leader, i.e. the assignment is still random.

Both models [M2] and [M5], lend credibility to the idea that more active individuals tend to get followed more by others, because of the higher likelihood of being observed by potential followers. Because Bechstein’s bats orientate via echolocation calls, potential followers coud detect and follow very active colony members even if those would not actively attempt to recruit followers via social calls (Fenton, 2003; Schöner *et al*., 2010; Knörnschild *et al*., 2012). This strengthen the position of [M5].

In conclusion, our investigations lend evidence to the hypotheses underlying [M5], namely that individuals have a natural tendency to follow, i.e. to acquire information socially from following a leader. Leaders, on the other hand, need not be detected because they actively recruit colony members (Schöner *et al*., 2010). They can be already sufficiently detected from observing their overall flight activity, when potential followers eavesdrop on the echolocation calls of potential leaders (Dechmann *et al*., 2009).

#### Identifying outliers in [M5]

As outlined in Section 2.4, null-models do not only allow us to narrow down the minimum complexity to reproduce some empirical findings. They also help us to identify subtle differences in the expected and observed networks. These cases indicate that the complexity in the model is insufficient in explaining observed behaviour and suggest that a more complex mechanism may be at play.

In our case, we notice four individuals - **1AC4** and **B1F6** (GB2 2008, Figure 6), **380C** (GB2 2011, Figure S1) and **A92B** (BS 2010, Figure S4) - whose centrality scores are considerably outside the 95% confidence range in model [M5]. Three of these four outliers are significantly more influential, i.e. have a larger centrality, than expected from [M5].

The mismatch means that they led more than expected from their flight activity alone. Since activity also includes behaviour unrelated to leading-following (e.g. individual exploration and revisits of boxes) and is normalized to the activity of others, it could be that these individuals actively engage in leading. This would give evidence to a *tendency to lead*, captured in model [M4], where we do not see the same outliers. This could indicate a more complex individual behaviour (e.g. specialisation in recruitment, higher recruitment efficiency or explicit preference for personal information gathering), as seen in honey bees (Seeley *et al*., 2006), which, otherwise, would have been difficult to single out.

Another reason for the mismatch in [M5] may be the number of followers that a leader has. A consistent tendency to have *several followers* per L/F event can result in centralities *larger* than expected from the rules underlying [M5]. But also the opposite can happen, as individual **B1F6** in the GB2 2008 data set illustrates. Its recruitment activity, as a leader in L/F events, is markedly *lower* than predicted based on its overall flight activity. In fact, most of its recordings in the data set came from discovery, exploration, and revisit events. Because the model uses the information about activity as an input, it predicts a higher centrality than observed in the real network. This discrepancy could indicate that the respective individual either does not attempt to spread the gathered information in recruitment events or did not manage to recruit followers.

#### Outlook

As suggested by Farine and Whitehead 2015, “there is a pressing need to combine network analysis with experimental manipulation.” Croft *et al*. 2011 highlights experimental manipulation as a very important tool to better understand social networks in animal populations. In the context of null models, this implies that proposed mechanisms could be used to design future empirical studies.

We believe that our results can contribute to this in two ways. First, our model candidates [M4] and [M5] for explaining recruitment need to be confirmed by field work. Experiments should possibly test whether recruitment is *passive* in the sense that followers randomly follow individuals that constantly need to echolocate for orientation and thus are detectable for potential followers. Alternatively, recruitment could be *active* in the sense that leaders intentionally attract the attention of their potential followers by e.g. acoustic signals or specific aerial displays. Both recruitment types have been suggested/demonstrated for other bat species (Fenton, 2003; Dechmann *et al*., 2009; Knörnschild *et al*., 2012; Schöner *et al*., 2010).

Second, the fact that we have identified specific individuals, whose recruitment behaviour cannot be explained by the given model complexity, allows to further investigate the characteristics of these individuals. For example, relating demographic, health or genetic characteristics to displayed inconsistencies with flight activity may reveal the reasons underlying their behavioural variability.

## Acknowledgements

We thank the local forestry and conservation departments for their continuous support throughout this long-term study, and the numerous people who helped gather the field data used in this study, in particular Anja Baigger and Markus Melber. This work profited strongly from the financial support of the German (DFG, KE 746/2-1/3-1/4-1/5-1/6-1) and the Swiss (SNF, 31-59556.99) national science foundations during the years covered in this study.

## S Electronic Supplementary Information

### S.1 Data set of leading-following events

Table S1 gives an example of the L/F data sets that were obtained in a previous study (CITE).

**Table S1:** Colony BS 2007, the L/F data are also used in Figure 2. `Leader_TS` and `Follower_TS` refer to the time stamps when the bats were recorded at the roost, `ID` refers to the roost ID. The last column `Info` tells us how the leader got informed about the roost. `personal` means she discovered it alone, `social` means she previously followed someone else.

| Leader_ID | Follower_ID | Leader_TS | Follower_TS | ID | Info |
| --- | --- | --- | --- | --- | --- |
| 0001F7D573 | 000644514C | 20070807T230329 | 20070807T230119 | 0b | personal |
| 000644514C | 0000624F85 | 20070807T231726 | 20070807T231734 | 0b | social |
| 0001F7D573 | 0000624F85 | 20070807T231457 | 20070807T231626 | 0b | personal |
| 0001F7D573 | 000697C596 | 20070807T231457 | 20070807T231939 | 0b | personal |
| 000644514C | 000697C596 | 20070807T231726 | 20070807T231939 | 0b | social |
| 00065EC1B3 | 0000624F85 | 20070807T232201 | 20070807T232008 | 0b | personal |
| 00065EC1B3 | 000697C596 | 20070807T232201 | 20070807T231940 | 0b | personal |
| 00065EC1B3 | 00068E1A46 | 20070807T232345 | 20070807T232344 | 0b | personal |
| 000697D368 | 00068E1A46 | 20070807T232758 | 20070807T232344 | 0b | personal |
| 00060F600D | 00068E1A46 | 20070807T232800 | 20070807T232344 | 0b | personal |
| 000697C596 | 00061791E7 | 20070807T235621 | 20070808T000117 | 0b | social |
| 00068E19F7 | 00061791E7 | 20070808T000406 | 20070808T000307 | 0b | personal |
| 00065EC1B3 | 00061791E7 | 20070808T000441 | 20070808T000307 | 0b | personal |
| 000697C596 | 000064BEA6 | 20070808T010145 | 20070808T010336 | 0b | social |

### S.2 Model-generated centralities

In addition to the Figures 3, 4, 5, 6 and 7 in the main text, Figures S1, S2, S3, S4 and S5 illustrate the model-generated vs. empirical centrality scores for all data sets in Table 1.

**Figure S1:**
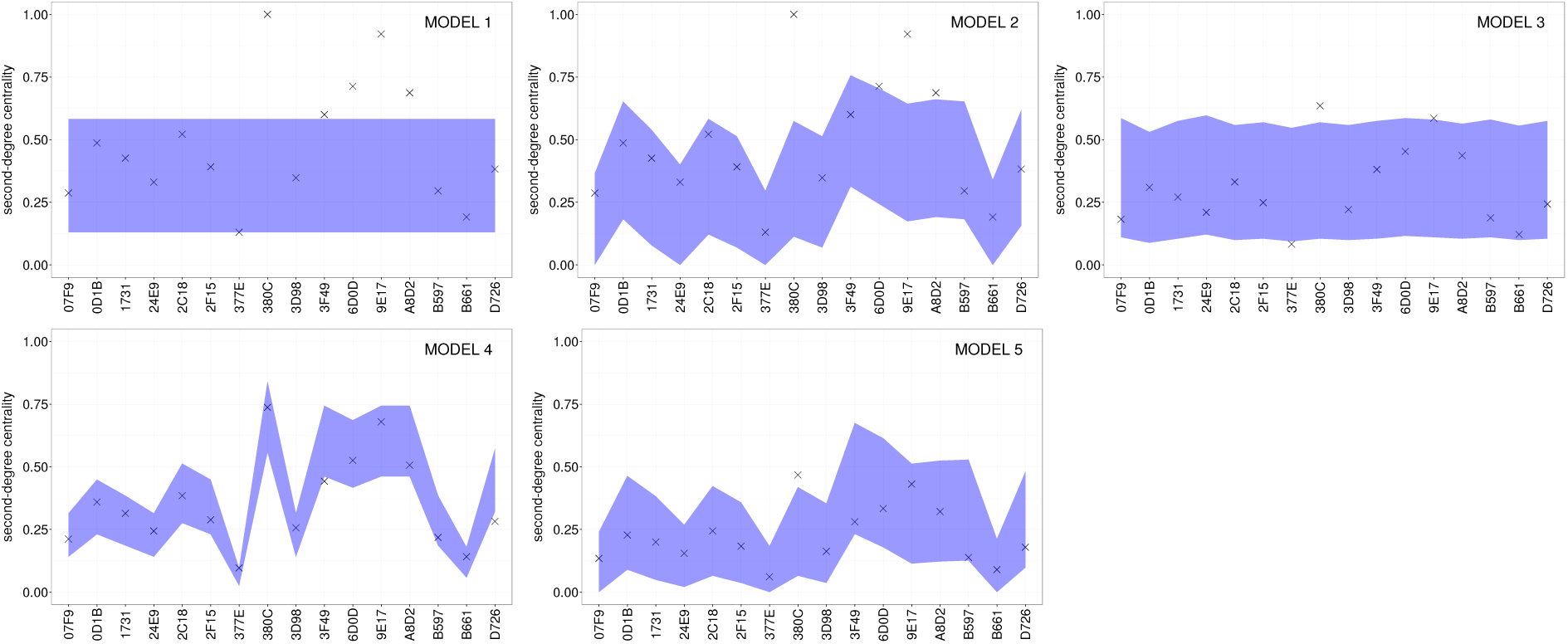
Model-generated vs. empirical centrality scores for colony GB2 in year 2011.

**Figure S2:**
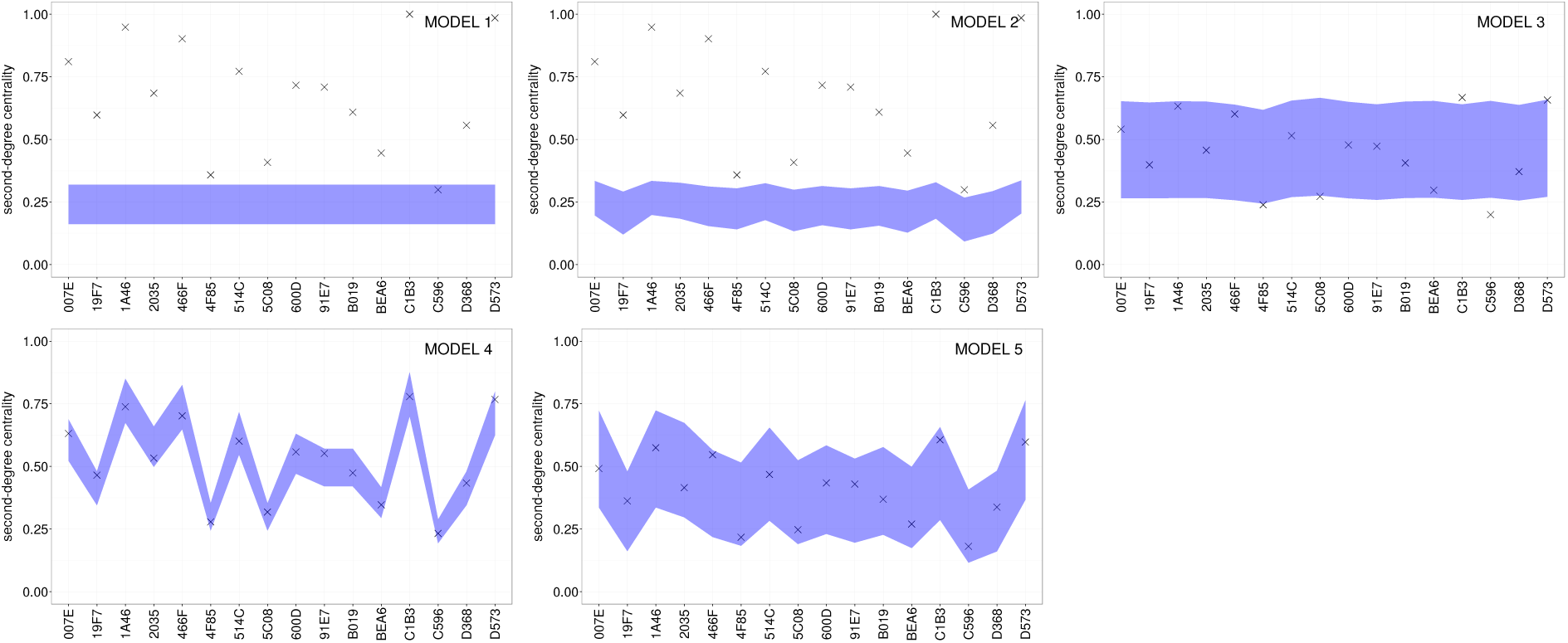
Model-generated vs. empirical centrality scores for colony BS in year 2007.

**Figure S3:**
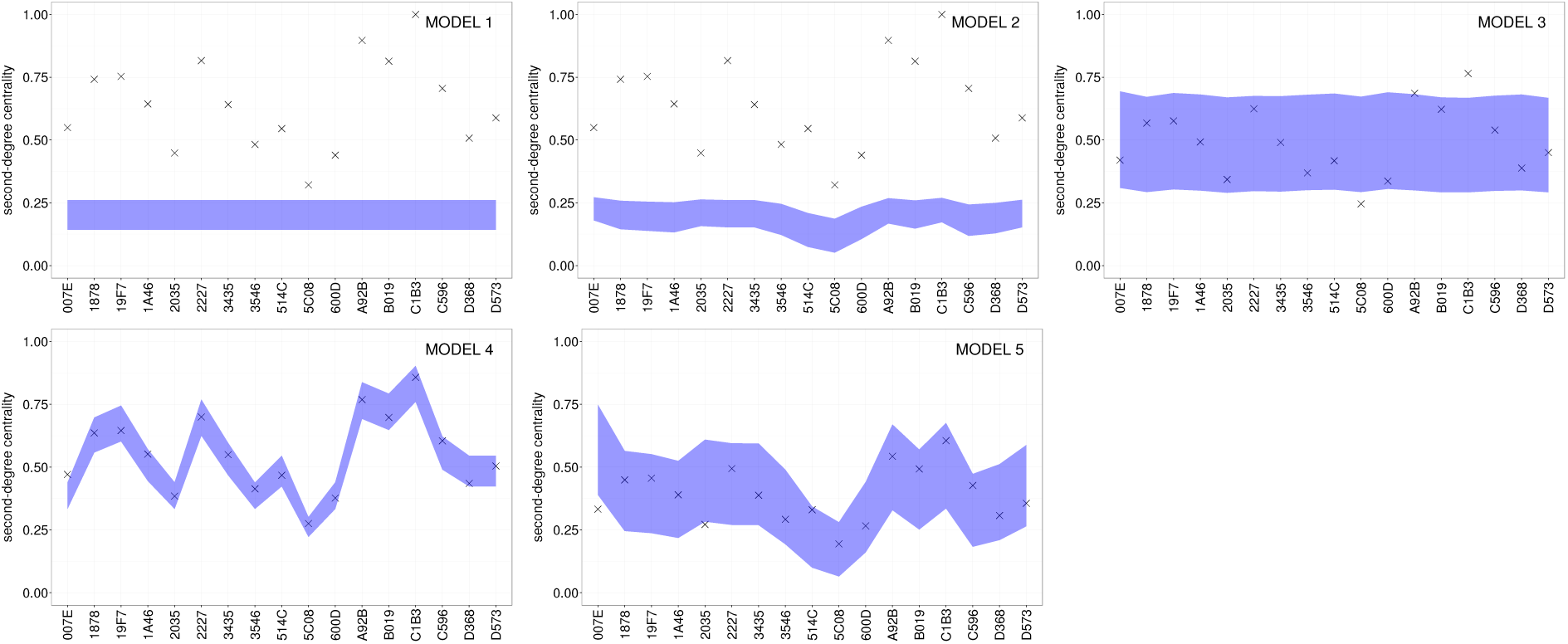
Model-generated vs. empirical centrality scores for colony BS in year 2009.

**Figure S4:**
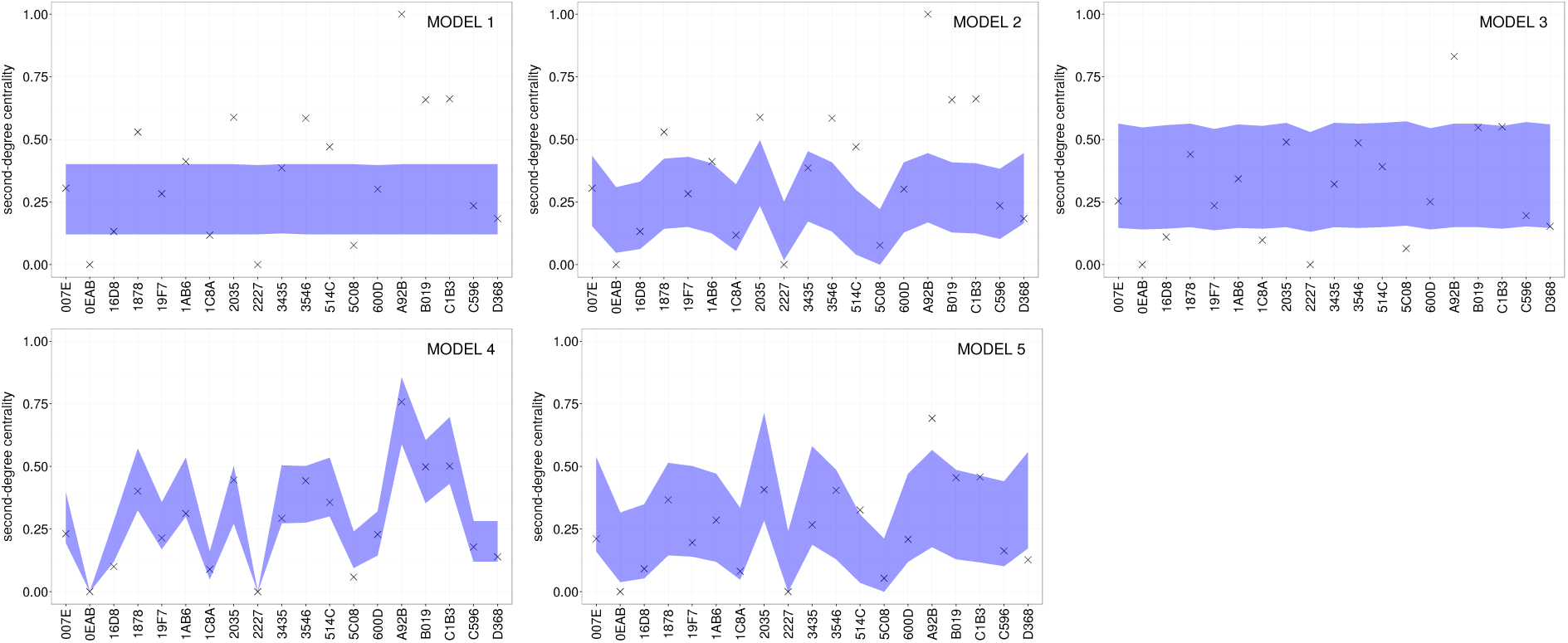
Model-generated vs. empirical centrality scores for colony BS in year 2010.

**Figure S5:**
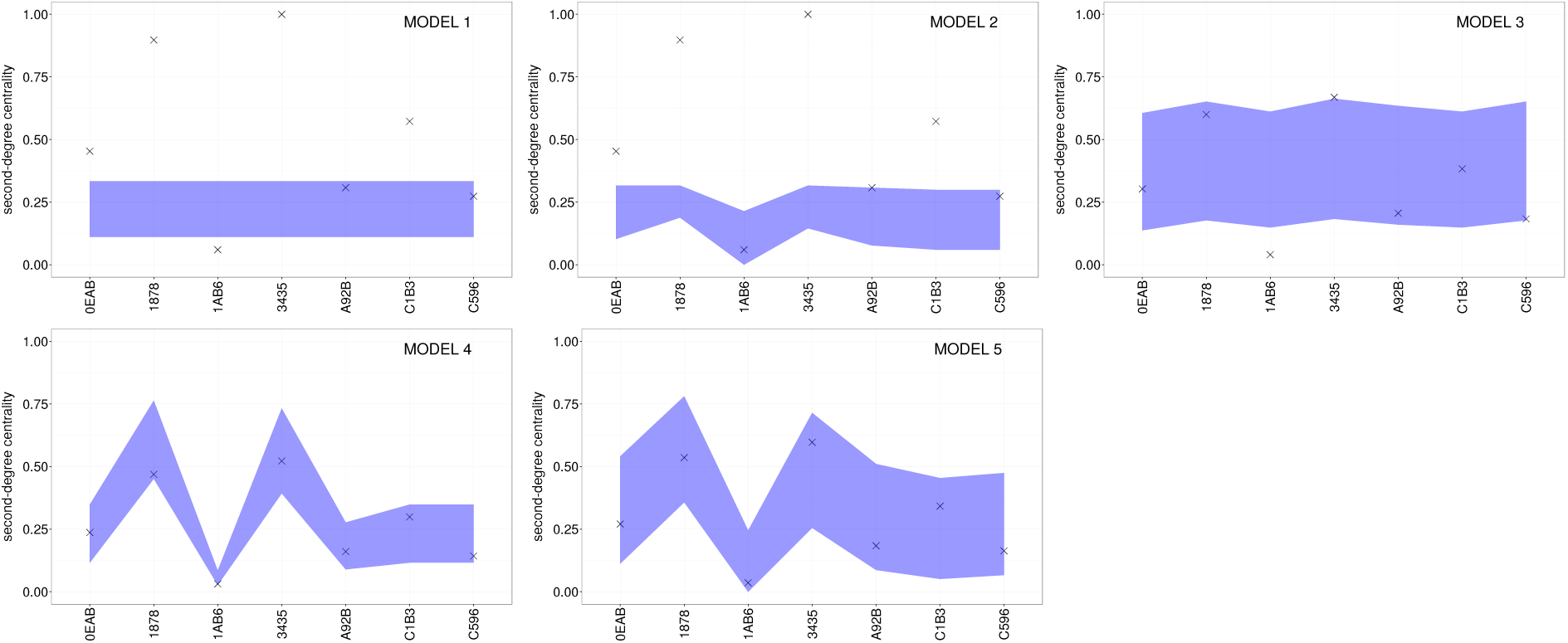
Model-generated vs. empirical centrality scores for colony BS in year 2011.

